# DreamDIA-XMBD: deep representation features improve the analysis of data-independent acquisition proteomics

**DOI:** 10.1101/2021.04.22.440949

**Authors:** Mingxuan Gao, Wenxian Yang, Chenxin Li, Yuqing Chang, Yachen Liu, Shun Wang, Qingzu He, Chuan-Qi Zhong, Jianwei Shuai, Rongshan Yu, Jiahuai Han

## Abstract

We developed DreamDIA-XMBD, a software suite for data-independent acquisition (DIA) data analysis. DreamDIA-XMBD adopts a data-driven strategy to capture comprehensive information from elution patterns of target peptides in DIA data and achieves considerable improvements on both identification and quantification performance compared with other state-of-the-art methods such as OpenSWATH, Skyline and DIA-NN. More specifically, in contrast to existing methods which use only 6 to 10 selected transitions from spectral library, DreamDIA-XMBD extracts additional features from dozens of theoretical elution profiles originated from different ions of each precursor using a deep representation network. To achieve higher coverage of target peptides without sacrificing specificity, the extracted features are further processed by non-linear discriminative models under the framework of positive-unlabeled learning with decoy peptides as affirmative negative controls. DreamDIA-XMBD is written in Python, and is publicly available at https://github.com/xmuyulab/Dream-DIA-XMBD for high coverage and precision DIA data analysis.

## 1 Introduction

Liquid chromatography coupled with tandem mass spectrometry (LC-MS/MS) has now become one of the most widely-used approaches for high-throughtput proteomic data acquisition due to its capability to quantify tens of thousands of peptides per hour^1, 2^. To meet the growing demand of large-scale quantitative proteome research, data-independent acquisition (DIA)^3–14^ mode was established. Instead of selecting specific precursors with higher intensities for fragmentation as in data-dependent acquisition (DDA), DIA encapsulated all precursors in a pre-designed isolation range for MS2 acquisition in an unbiased way^15^. It has been proven to outperform DDA and selected reaction monitoring (SRM) on various important aspects such as coverage, quantification accuracy and reproducibility^16–19^.

Despite various advantages of DIA, the challenge of DIA data analysis roots in its noisy spectra data originated from convoluted signals of multiple co-fragmented precursor ions. To overcome this problem, DIA data analysis is usually performed with the peptide-centric scoring (PCS) strategy^20^, where a spectral library that contains information of precursor ions of interests is queried against a series of raw data files to achieve higher sensitivity for large-scale complex biological samples^16, 21, 22^. In general, PCS software tools extract the elution profiles of the transitions in the library, identify and score them with a series of features and calculate the final discriminant scores for false positive control^23^. Naturally, both the extracted features and the discriminative models used to generate the final score determine, jointly, the performance of the software to produce the protein identification and quantification results. For the discriminative model, most DIA data analysis software tools use linear semi-supervised learning algorithms including semi-supervised linear discriminant analysis (ssLDA)^24^ and semi-supervised support vector machine (ssSVM)^25^. Non-linear discriminative models were also introduced in recent years, for instance the optional XGBoost classifier integrated in a new version of PyProphet^26^. DIA-NN^27^ also introduced a neural network-based model that gained great improvement. However, for the elution profile scoring methods, almost all the existing software tools use manually curated scoring systems based on expert knowledge such as Pearson correlations, shape of the elution profiles, relative intensities of the fragments, etc.^27–31^, which are heuristic and may not completely cover intrinsic characteristics of the complex elution patterns in DIA data. These heuristic feature extraction strategies may hinder the correct identification and accurate quantification of peptides and proteins in DIA projects.

Recently, deep representation learning has become a method of choice for feature extraction from unstructured data such as natural language^32^, speech^33^ and image^34^. Numeric results have confirmed that for these data, features from deep representation networks outperform those from conventional feature engineering strategies as the latter may not be able to exploit the intrinsic joint distribution of the signals in the high-dimension feature space towards the designated target of learning^35^. For DIA data analysis^36^, deep learning has also been recently introduced for the prediction of fragment intensities, retention time and ion mobility^37–41^, and de novo sequencing^42^.

In this paper, we introduce DreamDIA-XMBD, a PCS software suite for DIA data analysis based on the chromatogram features extracted by a deep representation network (Figure 1a, detailed descriptions of the data processing pipelines are in Methods). In contrast to most existing PCS software tools^29, 30^ which only consider 6 to 10 elution profiles for peptide scoring, DreamDIA-XMBD considers dozens of additional elution profiles for each precursor that has been ignored by most PCS software tools including all the theoretical fragment ions, potentially unfragmented precursor ions and isotopic peaks of each precursor (Figure 1b). These elution profiles are compiled into a set of representative spectral matrices (RSM), which are then further analyzed by a deep representation network based on the Long Short Term Memory (LSTM) model to capture more informative precursor features. The deep representation model in DreamDIA-XMBD has the capability to extract features from the complex elution patterns in RSM, which may bring only interference for conventional heuristic peptide scoring systems.

**Figure 1:**
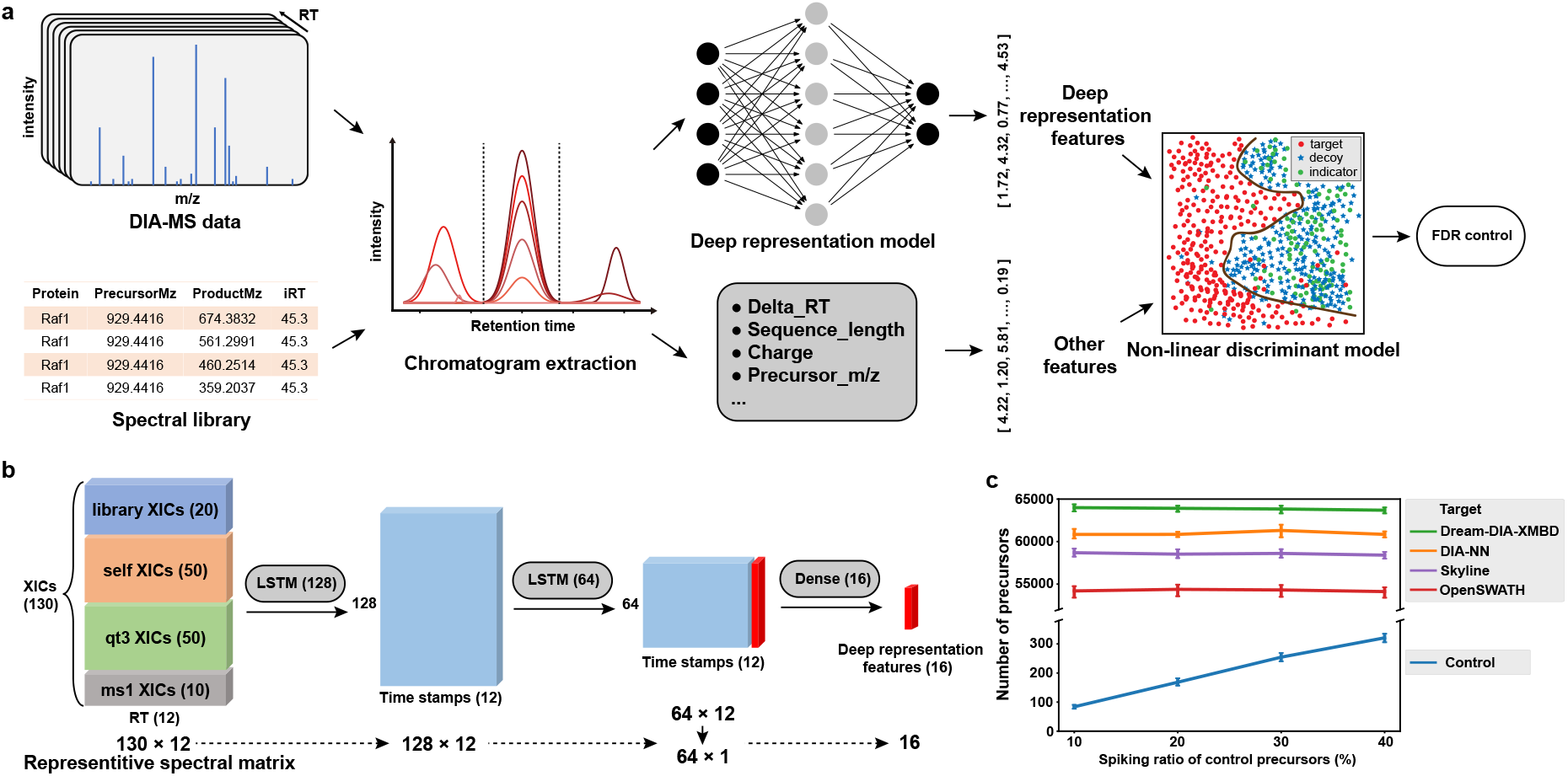
Schematic illustration of DreamDIA-XMBD and its identification performance. **a)** Schematic diagram of DreamDIA-XMBD. First, chromatograms of each target/decoy precursor and its corresponding fragment ions are extracted. Then the chromatograms are sent into the pre-trained deep representation model to obtain low-dimensional features. Subsequently, deep representation features combined with other features such as precursor m/z and charge are used for discriminant score calculation. **b)** Architecture of the deep representation model used by DreamDIA-XMBD. The input RSM contains four types of extracted ion chromatograms (XICs): fragment ions in the spectral library (library XICs); theoretical fragment ions of the precursor (self XICs); unfragmented precursor ion and its isotope peaks (qt3 XICs); precursor ion and its isotope peaks (ms1 XICs). The model contains two LSTM layers and a full-connection layer, by which the input RSM can be transformed to 16-dimension deep representation features. **c)** Identification performance of DreamDIA-XMBD on the mouse cerebellum dataset (MC dataset, acquired on Orbitrap Fusion Lumos mass spectometers, Thermo Fisher Scientific). Two-species spectral libraries are used for FDR control for fair comparison (Methods). For each spiking ratio of the control precursors in the spectral libraries, the numbers of target precursors (mouse) identified by different software tools under a fixed number of control (yeast or E.coli) precursors are plotted. Each point stands for the mean and deviation of the results from 10 parallel samples.

To fully untilize the deep representation features for precusor scoring, we further used XG-Boost as the discriminative model. Breifly, we trained an XGBoost classifier ba sed on all the precusors in the spectral library. Compared to linear discriminative models such as LDA where the decision boundary is limited to a linear hyperplane in the feature space, the XGBoost classifier has the advantage to enable non-linear decision boundary for better sensitivity. Moreover, to prevent overfitting, we followed the framework of positive-unlabeled learning^43^ by using the decoy precusors as affirmative negative controls in the training process to prevent the XGBoost classifer from picking up false targets (*π*_0_)^26, 44^ in the spectral library that are not detectable in a specific sample.

Finally, for accurate quantification, we calculated weighted area under the chromatogram for each fragment ion, where the weighting of each fragment is determined as the sum of its Pearson correlations with all the other fragments from the same precursor based on the hypothesis that noise caused by coeluted peptides should have low correlations with the other true chromatograms.

We compared the identification and quantification performance of DreamDIA-XMBD with several state-of-the-art open-source PCS software tools including OpenSWATH^29^, Skyline^30^ and DIA-NN^27^. Our proposed method outperformed the other tools with more target precursors identified in the two-species library test^45^ and more accurate quantification results in the LFQbench test^46^. DreamDIA-XMBD provides a deep representation network based feature extraction method for DIA data analysis, in combination with a novel interface for deep learning algorithms to be introduced to obtain better performance for large-scale biological and medical proteome research. The training data of the deep representation model used in DreamDIA-XMBD can be easily obtained from public datasets. We have provided two trained models that can be directly applied to analyze DIA data, as well as an application programming interface (API) for customized model. DreamDIA-XMBD is publicly available at https://github.com/xmuyulab/Dream-DIA-XMBD.

## 2 Methods

### Deep representation models in DreamDIA-XMBD

The key step of the DreamDIA-XMBD algorithm is to extract relevant features of the chromatograms with deep representation models. The input of the deep representation models is RSM (Figure 1b), a matrix consisting of 130 top elution profiles selected based on their intensities from four types of elution profiles including *library*, *self*, *qt3* and *ms1*. The *library* part contains 20 top XICs of fragment ions in the spectral library. The *self* part contains 50 top XICs of all theoretical fragment ions of each precursor. For precursors with two charges, fragment ions with one charge are considered. For precursors with charges greater than two, fragment ions with one and two charge(s) are considered.. The *qt3* part^47^ contains top 50 XICs of unfragmented precursor and its isotope peaks in MS2 spectra. The *ms1* part contains top 10 XICs of precursor itself and its isotope peaks. The retention time width of RSM is set to 12 cycles by default, which was long enough for most elution signals by our visual verification on several DIA datasets.

The deep representation network used in this work consists of two LSTM layers and two full-connection layers. The first LSTM layer has 128 neurons with input dropout of 0.4 and recurrent dropout of 0.3. The second LSTM layer with 64 neurons and the same dropout settings was then stacked on the first layer. Two full-connection layers with 16 and 1 neuron(s) respectively were added on the top of the model. Rectified linear unit (ReLU) activation function was used for hidden layers, while sigmoid function was used for the final layer to obtain an output ranging from 0 to 1. All the deep representation models were built with Keras in Python. The models were trained on the RSMs in the training datasets as a binary classifier to differentiate real precursors and decoys with a cross-entropy loss function. The output of the final layer of the trained deep representation model, which hereafter referred to as the deep discriminant score (*dds*) as it represents the likelihood that a certain RSM is from a target peptide present in the sample as seen by the network, is used for RT normalization and peak picking in DreamDIA-XMBD. In addition, the 16-dimension output of the second to last layer of the model is used as the deep representation feature for the discriminative model to produce the final discriminant scores for each precusor from the spectral library (Figure 1b).

### Building sample-specific spectral libraries

The raw data files were first transformed to centroided mzXML files by ProteoWizard^48^ (version: 3.0.19317) with all the other arguments unchecked. The resulting mzXML files were processed by DIA-Umpire to generate pseudo MS/MS spectra. X!Tandem^49^ and Comet^50^ were used to search these pseudo spectra. The searching results were then filtered by PeptideProphet^51^ and ProteinProphet^52^ in TPP^53, 54^. Finally, the sample-specific spectral libraries were generated by SpectraST^55^.

### Training data preprocessing of the deep representation models

The training data of the deep representation model of DreamDIA-XMBD (1.0.0) were generated as follows. First, sample-specific spectral libraries were built as stated earlier. The resulting spectral libraries were further processed by DreamDIA-XMBD to generate decoys. Subsequently, OpenSWATH and Pyprophet^26^ were used to find RT of each target or decoy precursor in the spectral libraries across all runs. All the necessary XICs were extracted and saved as RSMs. In total two deep representation models were trained. A generic model was trained based on data from whole cell lysates of HEK 293 cells^56^ from xxx. In addition, a special model from SCIEX TripleTOF 5600 was trained on data were from the L929 mouse dataset^57^ as data from it has different characteristics compared with those of other spectrometers (Supplementary Note 3).

### DreamDIA-XMBD workflow

DreamDIA-XMBD supports centroided .mzML or .mzXML MS data files as input. In addition, it integrates the cross-platform MS file conversion tool ThermoRawFileParser^58^ to read .raw files directly from Thermo Fisher equipments. After reading a spectral library, DreamDIA-XMBD generates a decoy for each precursor ion by random shuffling of the sequence using Fisher-Yates algorithm. Decoys generated from other software tools for instance OpenSWATH can also be used by DreamDIA-XMBD.

After acquiring the necessary inputs file, DreamDIA-XMBD randomly samples a set of endogenous precursors in the library for RT normalization. DreamDIA-XMBD searches for the best RSM based on the value of *dds* provided by the deep representation model for each precursor in the sampled set across the whole retention time gradient. Then it fits a linear model based on time points of these best RSMs against their normalized RT, by which the RTs of the other precursors in the spectral library can be predicted. The Random Sample Consensus (RANSAC)^59^ algorithm is used to detect outliers. Subsequently, DreamDIA-XMBD extracts RSMs within a predefined range centered on the predicted RT for each target or decoy precursor in the spectral library, and keep the RSMs with *dds* higher than a cut-off value.

To distinguish between target and decoy precursors, DreamDIA-XMBD fits a non-linear discriminative model based on the deep representation features in combination with other features such as peptide lengths and charges (Supplementary Note 1). Due to the existence of false targets in a spectral library, the discriminative model is trained based on the principle of positive-unlabeled learning^43^ with decoy peptides as affirmative negative controls while treating targets as unlabeled.

XGBoost was chosen as the default discriminative model for its superior performance (Results). However, the software also provides the option for users to choose other classifiers such as random forest. To furhter overcome the inaccuracy of RT prediction, we adopt the test-time augmentation strategy where all the RSMs with higher scores given by the deep representation model are considered as evidences for a precursor.

After discriminant score calculation, the RSMs with the highest discriminant score for each precursor are kept for FDR control. DreamDIA-XMBD estimates FDR by dividing the number of target precursors by the number of all precursors with discriminant scores exceeding a cut-off score. For a specified FDR level, DreamDIA-XMBD can search for a proper cut-off score for valid identifications. Protein-level FDR is calculated similarly through dividing the number of target precursors by the number of all precursors exceeding a cut-off score for each protein respectively.

### Peptide and protein quantification

For peptide quantification, DreamDIA-XMBD uses a weighted area method to prevent the influence brought by other co-eluting ions as follows.

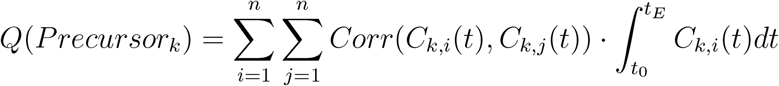

Herein, *C*(*t*) means elution chromatogram of an ion. *Corr*() is the Pearson correlation of two fragment ions, and the integration term calculates the area under the ion’s chromatogram. DreamDIA-XMBD quantifies all fragment ions according to the areas of their chromatograms for each precursor. The weight of each fragment is calculated as the sum of the Pearson correlations with the other fragments, and the quantification result of a precursor is the weighted sum of the top six fragments associated with it. For protein level quantification, DreamDIA-XMBD sums the intensities of the top three abundant precursors for each protein.

### Two-species spectral library method

As different software tools have different strategies for FDR calculation, it is difficult to directly compare the identification performance according to their results. Therefore, we adopt the two-species spectral library method that had been used in previous work for benchmarking^19, 27, 45^, where different ratios of the proteins from another different species were added to the sample-specific spectral libraries as target precursors, which could then be used to evaluate the false positive identification of an algorithm.

### Software versions for comparison

DreamDIA-XMBD (1.0.0) was compared with OpenSWATH^29^ (2.6.0), Skyline^30^ (19.1.0.193) and DIA-NN^27^ (1.7.11).

### Benchmarking of precursor identification

The mouse MC dataset, SGS human dataset and HeLa dataset were first processed to produce sample-specific libraries. Peptides that belonged to multiple proteins were discarded. Then we extracted yeast and E.coli precursors, which were not expected to exist in the sample-specific libraries, as the control precursors from the mixed library built by LFQbench^46^. These precursors were filtered to discard sequences that existed in the sample-specific libraries. Next, we spiked them into the sample-specific libraries with different ratios to build the two-species libraries. Due to different sizes of the sample-specific libraries and the number of control precursors of different datasets, the ratios of the two-species library were ranging from 10% - 40%, 10% - 200% and 10% - 50% for the MC dataset, the SGS human dataset and the HeLa dataset, respectively.

Top 6 fragment ions with the highest intensities for each precursor were retained. Subsequently, all the sample files were processed by the software tools with the resulting two-species libraries. For DreamDIA-XMBD, centroided mzXML files were used as input. For Skyline and DIA-NN, centroided mzML files were used. For OpenSWATH, profile mzXML files were used as recommended^60^. After analysis, the identification results were compared at the same control levels. For fair comparison, we filtered results from all software tools by their discriminant scores to have the same number of control precursors, where the number is set to FDR of 1% by DIA-NN (Figure 1c, Supplementary Figure S1, Supplementary Figure S2).

### Configuration of the software tools

For DreamDIA-XMBD, default settings were used except that the “--swath” option was specified for SCIEX TripleTOF 5600 data analysis. For DIA-NN, the Linux commond-line tool with default settings was used. OpenSWATH was run with options “-readOptions cacheWorkingInMemory -batchSize 0 -rt extraction window 1200 -threads 20”. Then the output was processed by PyProphet-cli (0.0.19)^26^ with “--lambda=0.4 --statistics-mode=local” options. Suboptimal peak groups were subsequently discarded. For Skyline, the step-by-step settings are described in Supplementary Note 2. We did not perform extensive parameter optimization to obtain best results for Skyline and OpenSWATH, as it had already been proved^27^ that DIA-NN performs better than both of them, which is also proven by our results.

## 3 Results

### DreamDIA-XMBD shows better identification performance

We benchmarked DreamDIA-XMBD against several mainstream open-source PCS software tools on various public datasets. First, the performance of peptide identification was tested on the mouse cerebellum dataset^61^ (MC dataset, acquired on Orbitrap Fusion Lumos mass spectometers, Thermo Fisher Scientific) and the SWATH-MS Gold Standard human dataset^29^ (SGS human dataset, acquired on TripleTOF 5600 System, SCIEX). All data were processed by OpenSWATH^29^, Skyline^30^, DIA-NN^27^ and DreamDIA-XMBD separately. We used the two-species spectral library method, which has been used for unbiased software benchmarking^19, 27, 45^, for our comparisons (Methods). The results show that DreamDIA-XMBD achieves the best identification performance compared with the other software tools (Figure 1c and Supplementary Figure S1). For data generated from either Thermo Fisher DIA (Figure 1c) or SWATH (Supplementary Figure S1) equipments, DreamDIA-XMBD identified more target precursor ions under the same control levels. We also benchmarked DreamDIA-XMBD on the HeLa dataset (acquired on QExactive HF, Thermo Fisher Scientific) used in DIA-NN^27^ paper (Supplementary Figure S2). DreamDIA-XMBD still achieved the best identification performance compared with the other software tools for data acquired at different gradient lengths.

### DreamDIA-XMBD produces reliable improvements

Although we have filtered the sample-specific libraries in a relatively strict approach, we cannot regard these target precursors as absolute ground truth for identification benchmarking. To better illustrate the reliability of the improvements brought by DreamDIA-XMBD, we introduced high confident proteins (HCPs), which were defined as proteins with more than 100 identified precursors in at least one run for all the software tools. We then monitored the numbers of their distinctive precursors reported by different software tools. Theoretically, precursors represent evidences for each protein to be identified, thus the more precursors reported, the higher chance for a protein to exist in the sample. As for HCPs, more precursors reported indicates higher reliability of the software. We found that DreamDIA-XMBD could identify more precursors for most HCPs compared with the other software tools (22, 20 and 27 out of 29 HCPs compared with DIA-NN, Skyline and OpenSWATH respectively), which indicated that the improvement brought by DreamDIA-XMBD was reliable (Figure 2).

**Figure 2:**
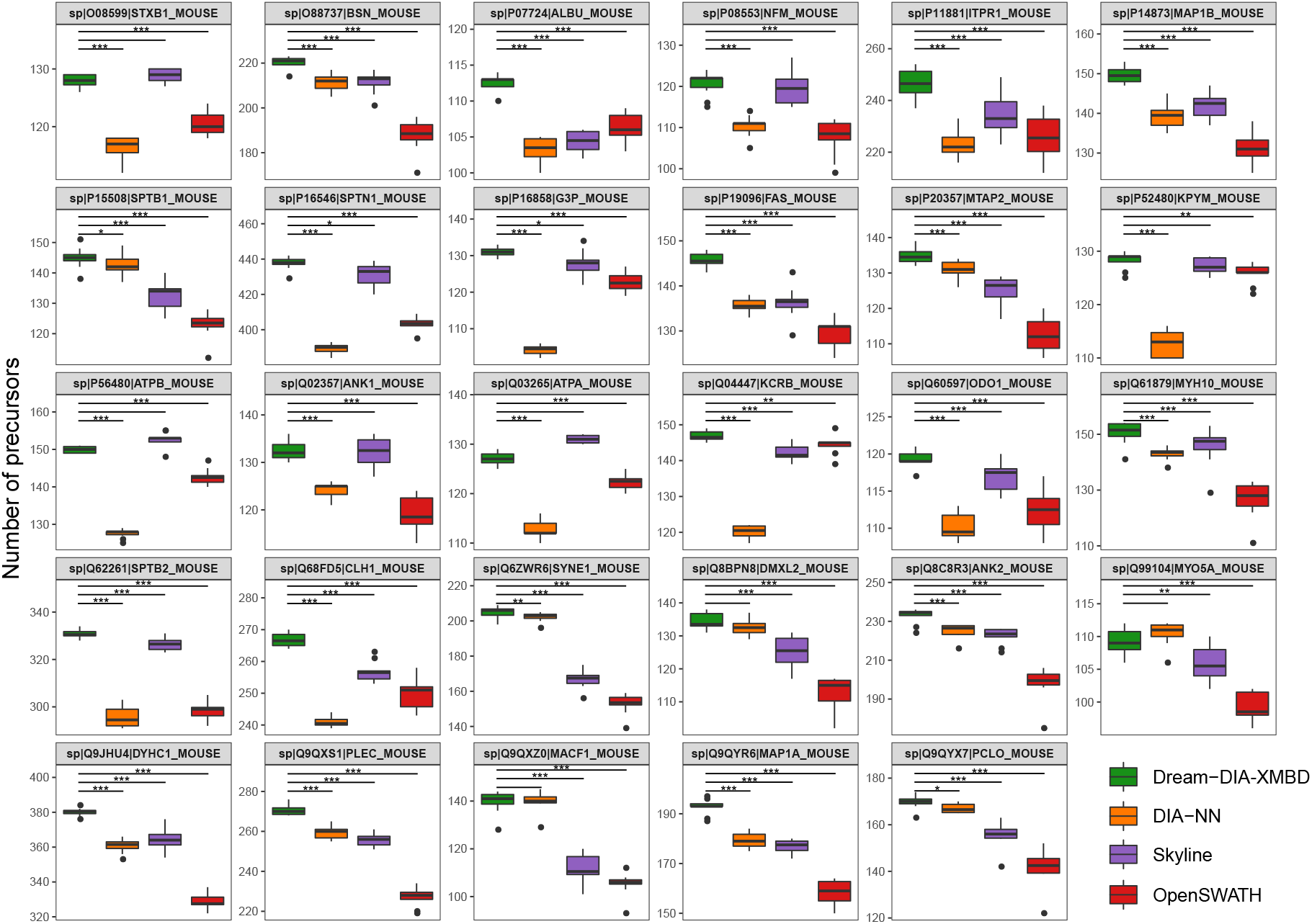
Boxplots of the numbers of precursors for 29 HCPs reported by different software tools on the MC dataset. Each box stands for the distribution of the numbers of target precursors identified for each HCP across 10 runs. Spectral library with 40% spiking ratio of control pre-cursors was used. Single-sided Wilcoxon rank sum tests were performed between the results of DreamDIA-XMBD and the other three software tools. The numbers of “*” denote the statistical significance of the tests (*: p-value*<*0.05; **: p-value*<*0.01; ***: p-value*<*0.005). Among the 29 HCPs, DreamDIA-XMBD reported significantly more precursors for 22, 20 and 27 HCPs when p-value*<*0.005, and 24, 21 and 29 when p-value*<*0.01 respectively compared with DIA-NN, Skyline and OpenSWATH.

### Deep representation models can extract peptide relevant information from chromatograms

What makes DreamDIA-XMBD a better peptide identification method is that the deep representation models it uses extract more relevant information from the chromatograms. Compared with expert knowledge, the deep representation models take advantages of previously acquired data to understand the intrinsic properties of elution pattern that represent a real peptide in a more precise and stable manner. We find that positive and negative precursors’ RSM can be visually separated on the t-SNE dimension reduction embedding of the 16-dimension deep representation features (Supplementary Figure S3), which indicates that these features are highly informative regarding positive and negative precursors, and therefore contribute to better identification performance under the same FDR compared with traditional methods.

### Non-linear discriminative model improves the performance of DreamDIA-XMBD

The final discriminant score of each precursor in DreamDIA-XMBD is calculated by a binary classifier instead of direct output from the deep learning model itself (Figure 1a). This strategy not only enables extra features that cannot be described by the deep learning model to be involved in the identification step (Figure 1a), but also improve the generalization capability of the algorithm. Moreover, as indicated by the t-SNE map, the distribution of the precursors in the feature space can be highly non-linear (Supplementary Figure S3), which is difficult for linear classifiers to discriminate between real peptide signals and noise. In such case, a non-linear classifier could obtain better identification results. To validate this hypothesis, we tested 7 commonly-used machine learning algorithms including LDA, logistic regression, CART decision tree, Adaboost, gradient boosting decision tree (GBDT), random forest (RF) and XGBoost on the MC dataset (Figure 3). Among all the tested classifiers, tree-based ensemble models including gradient boosting decision tree (GBDT), XGBoost and random forest (RF) obtained better performance due to their non-linear decision boundary, while linear models such as logistic regression and LDA in general performed worse than other non-linear models. XGBoost achieves the best performance, and is chosen as the default discriminative model in DreamDIA-XMBD.

**Figure 3:**
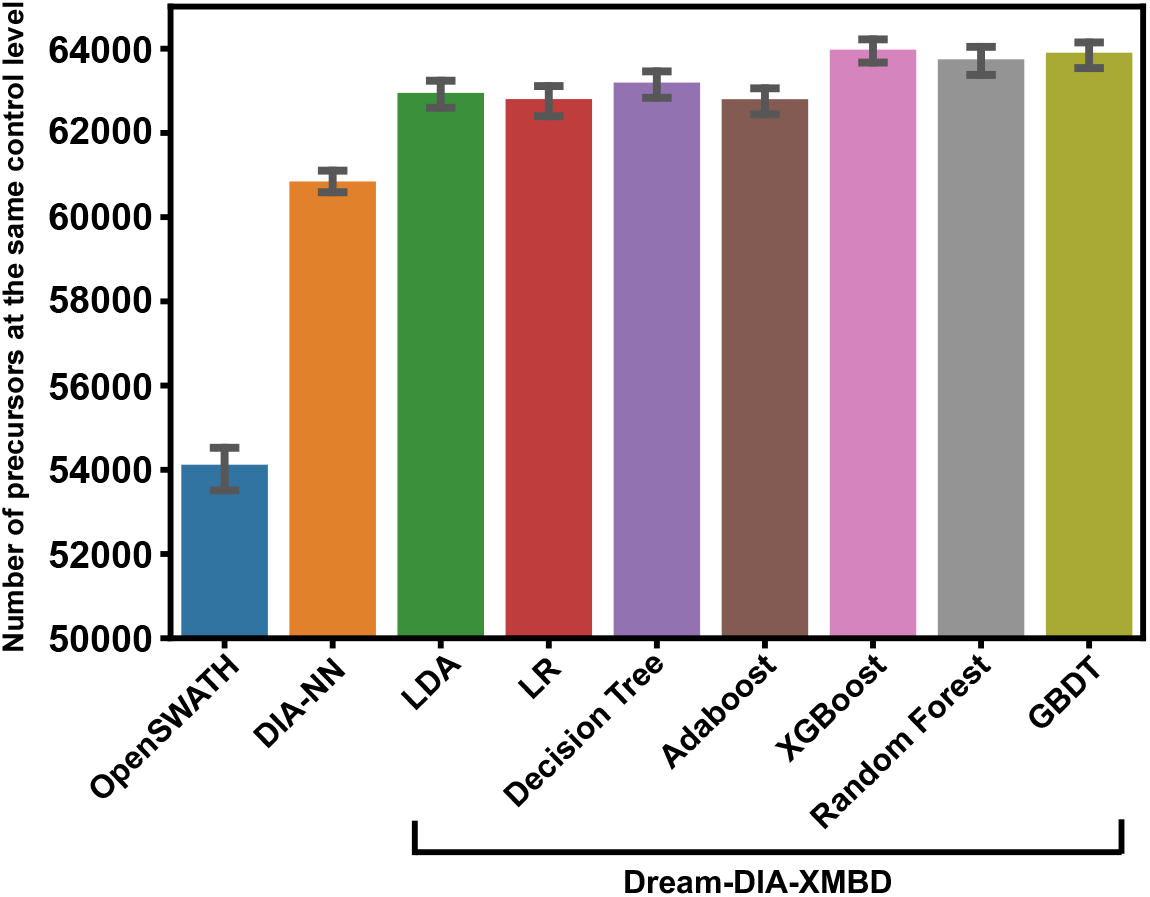
Comparison of identification performance of various discriminative models on the MC dataset. The spiking ratio of control precursors is 40%. The numbers of target precursors (mouse) are shown at the same control level from 1% FDR given by DIA-NN.

### DreamDIA-XMBD has better quantification performance

We benchmarked quantification performance of DreamDIA-XMBD against OpenSWATH and DIA-NN using LFQbench software suite^46^. The internal dataset of LFQbench contains known ratios of the proteins from different species (human, yeast and E.coli), which can be used to evaluate the accuracy and stability of a quantification algorithm by monitoring the recovery levels of these ground truth ratios. We chose the HYE124 samples with 64-window setup acquired from TripleTOF 6600 systems for quantification performance benchmarking. Each software tool should output results at 1% FDR based on its own standard. We kept precursors that had been reported at least once by all the three software tools. In total, 16556 human precursors, 11892 yeast precursors and 10390 E.coli precursors were retained for comparison. Compared with OpenSWATH and DIA-NN, DreamDIA-XMBD shows better quantification performance for the peptides and proteins of yeast and E.coli (Figure 4).

**Figure 4:**
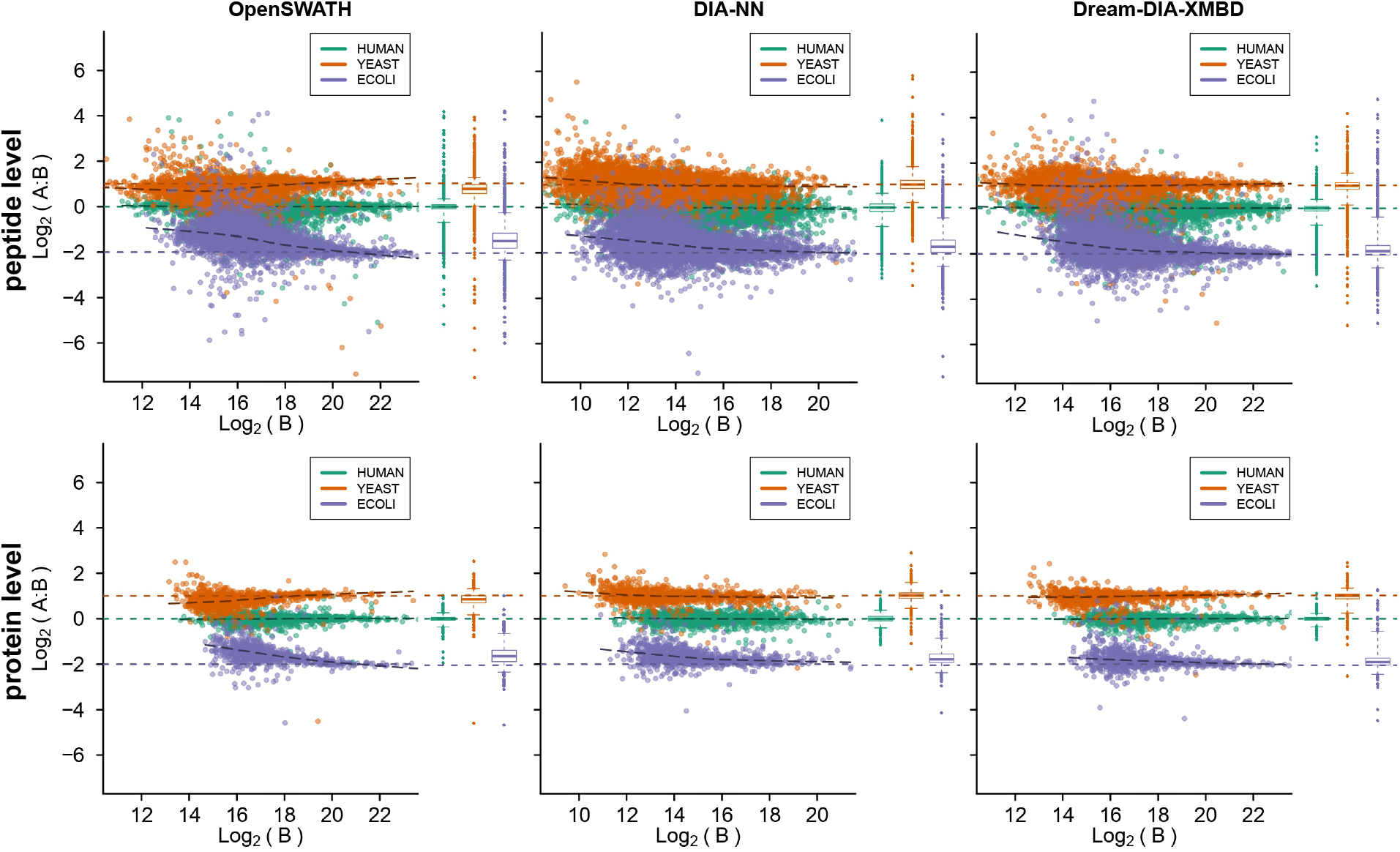
Quantification performance evaluation with LFQbench on the HYE124 datasets. Quantification results of peptides (the first row) and proteins (the second row) from OpenSWATH, DIA-NN and DreamDIA-XMBD. All results were obtained at 1% FDR for each sample, and 16556 human precursors, 11892 yeast precursors and 10390 E.coli precursors that reported by all the three software tools in at least one sample were analyzed and compared.

## 4 Discussion

We developed DreamDIA-XMBD, a deep nerual network based DIA data analysis software. By adopting a data-driven approach, the deep representation network used in DreamDIA-XMBD learns a more comprehensive representation of the distribution of the highly-variable elution profiles of peptides generated by DIA, thus improves the identification and quantification performance at both peptide and protein levels. DreamDIA-XMBD consists of a complete DIA data analysis pipeline from raw acquired data to quantification results and can be independently used without assistance of any other software. It is anticipated that with the increasing amount of DIA data for knowledge extraction and model training, deep learning based data-driven strategy can be a preferred method to overcome the limitations of DIA to further improve its coverage and precision for large-scale DIA proteome research. The RSM data structure and programming framework introduced by DreamDIA-XMBD can be a start towards this goal.

## 5 Key points

We developed DreamDIA-XMBD, a DIA data analysis software tool that introduced deep representation networks to extract informative features from elution profiles in DIA data in place of conventional heuristic peptide scoring systems.

DreamDIA-XMBD shows better peptide identification and quantification performance compared with several state-of-the-art software tools including OpenSWATH, Skyline and DIA-NN.

DreamDIA-XMBD provides a universal data structure and framework for further introduction of deep learning methods to obtain better performance for large-scale biological and medical proteome research in the future.

## Data Availability

The SGS dataset is available at PeptideAtlas raw data repository with accession number PASS00289. Other data used in this study were also public datasets deposited to the ProteomeXchange Consortium via the PRIDE^62^ or iProX^63^ partner repository. The dataset identifiers include PXD015098, PXD021390, PXD011691, PXD005573 and PXD002952. The source data of all the figures in this paper are available in supplementary materials.

## Code Availability

DreamDIA-XMBD (1.0.0) is open-source and available at https://github.com/xmuyulab/Dream-DIA-XMBD.

## Author Contributions

M.G. and R.Y. designed the study and the algorithms. M.G., W.Y. and R.Y. wrote the first manuscript. M.G., W.Y. and S.W. implemented the algorithms. M.G., C.L., Y.C., Y.L., Q.H. and C.Q.Z performed the experiments. J.S. and J.H. gave advices on the algorithm and experiment designs. All authors discussed and commented on the manuscript.

