## Supplementary for "DreamDIA-XMBD: deep representation features improve the analysis of data-independent acquisition proteomics"

Gao et al

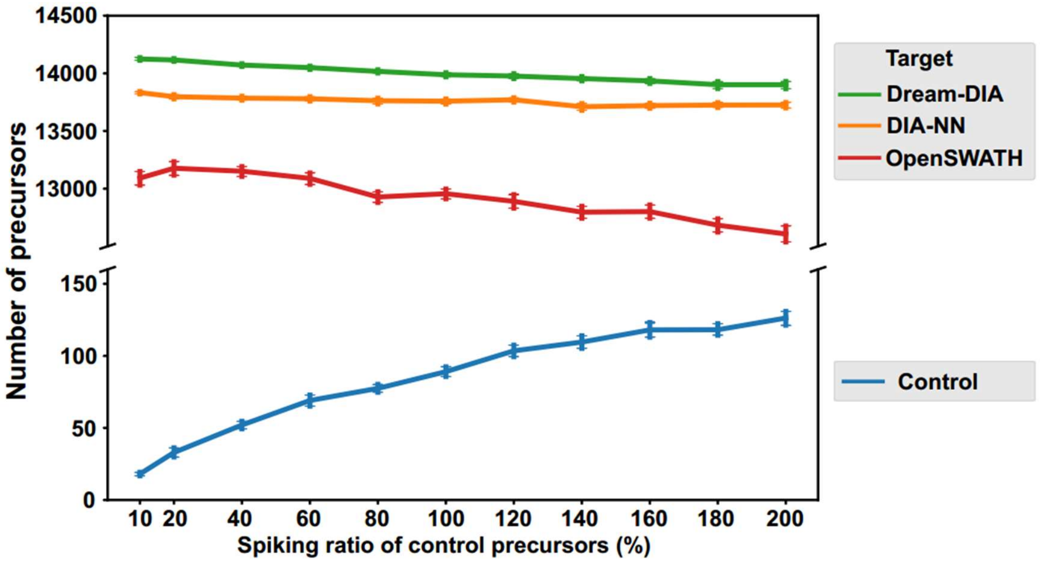


**Supplementary Figure S1.** Identification performance of DreamDIA-XMBD on the SWATH-MS Gold Standard dataset (SGS dataset, acquired on TripleTOF 5600 System, SCIEX). For each spiking ratio of the control precursors in the spectral libraries, the numbers of target (human) precursors identified by different software tools under a fixed number of control (yeast or E.coli) precursors are plotted. Each point stands for the mean and deviation of the results of 30 parallel samples in the SGS dataset.

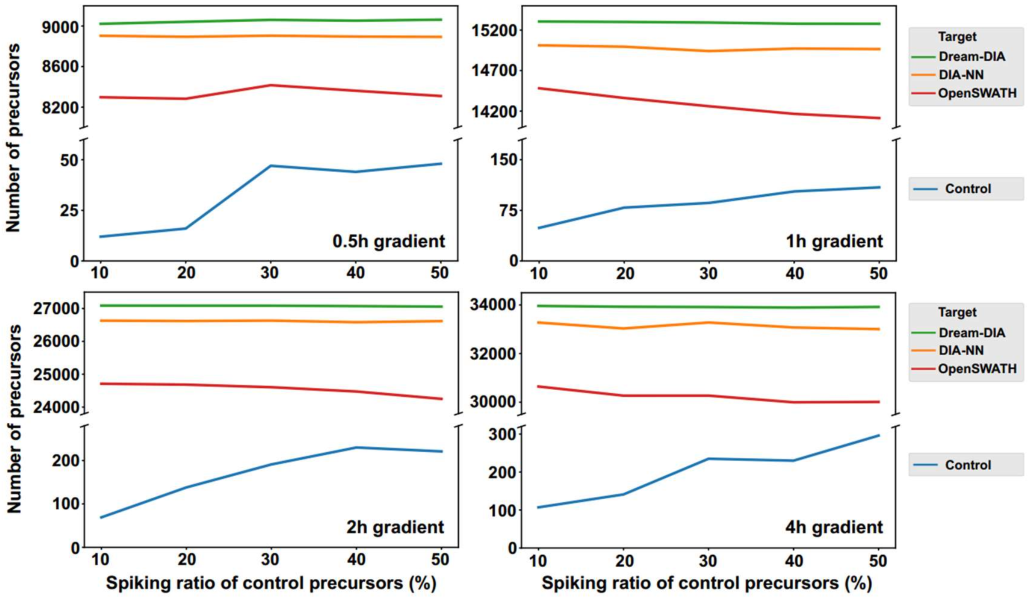


**Supplementary Figure S2.** Benchmark of identification performance of DreamDIA-XMBD on the HeLa dataset (acquired on QExactive HF, Thermo Fisher Scientific) used by DIA-NN. Different gradient lengths from 0.5h to 4h were tested. For each spiking ratio of the control precursors in the spectral libraries, the numbers of target (human) precursors identified by different software tools under a fixed number of control (yeast or E.coli) precursors are plotted.

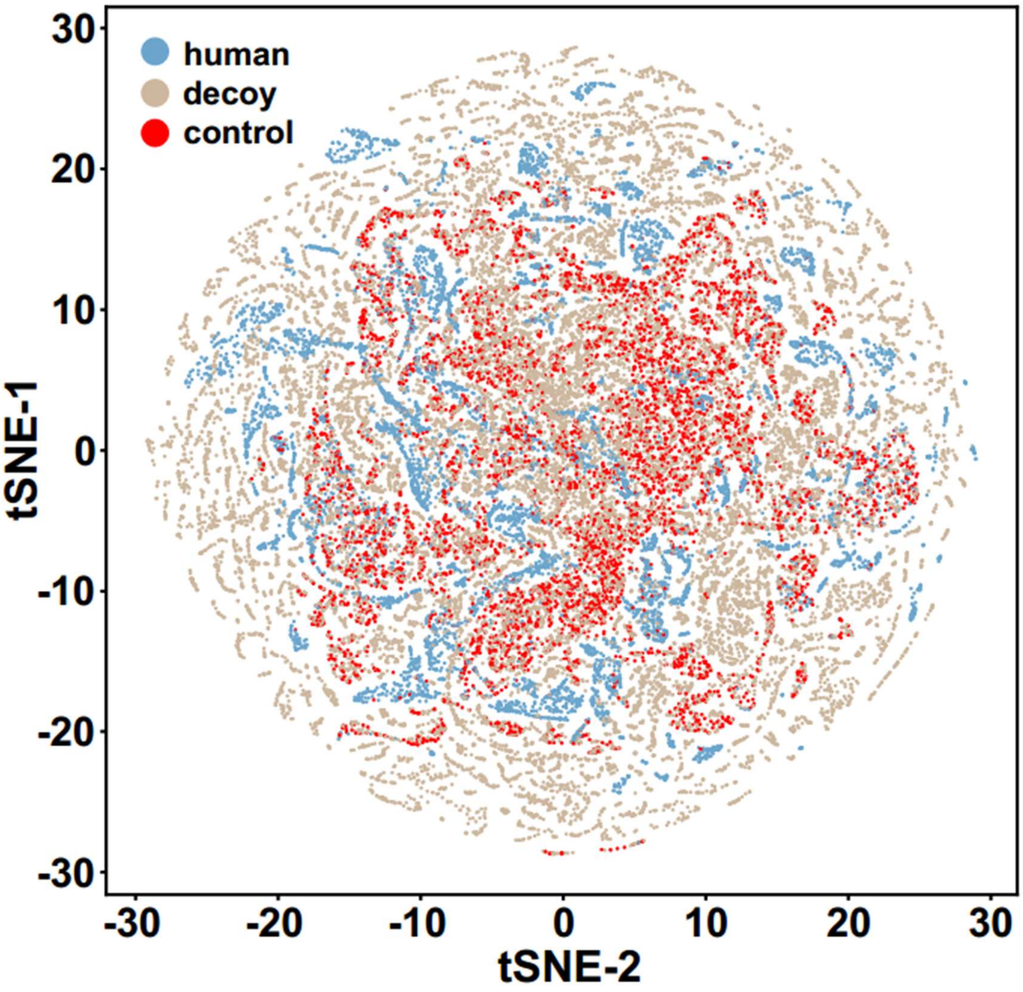


**Supplementary Figure S3.** tSNE of the extracted 16-dimension deep representation features by DreamDIA-XMBD from a sample of SGS dataset. Two-species library was used in which the spiking ratio of control precursors was 200%. Each point stands for a candidate peak group of a precursor. Only 5% of the extracted peak groups are displayed here for better visual impression (human: 7190; decoy: 21316; control: 5564).

### Supplementary Notes

#### 1. Auxiliary scores in DreamDIA-XMBD

Several auxiliary scores that cannot be contained in the deep representation models for candidate peak groups of each precursor in DreamDIA-XMBD are listed below.

(1) Difference between the real RT and the RT recorded in the library.
(2) Square of the difference between the real RT and the RT recorded in the library.
(3) Cosine similarity of the real intensities and library intensities of all the fragments.
(4) Mean and standard deviation of the three scores above of all the candidate peak groups for each precursor.
(5) Length of the peptide sequence.
(6) Charge of the precursor.
(7) m/z of the precursor.

#### 2. Skyline step-by-step settings

We analyzed the MC data by Skyline [1] for the performance benchmarking of DreamDIA-XMBD. Actually, we followed most settings provided by LFQbench [2], and the detailed procedures were shown below.

(1) Open Skyline.
(2) Import transition list.
(3) Close the "Settings" window.
(4) Settings -> Transition Settings:

- Full-Scan:
  - Acquisition method: DIA;
  - Product mass analyzer: TOF;
  - Isolation scheme: input the isolation window settings manually;
  - Resolving power: 100,000;
  - Retention time filtering: Use only scans within 10;
- Instrument:
  - Min m/z: 50 m/z;
  - Max m/z: 2000 m/z;
  - Method match tolerance m/z: 0.01 m/z;
- Library:
  - Ion match tolerance: 0.5 m/z;
  - If a library spectrum is available, pick its most intense ions: checked;
  - Pick: 6 product ions
  - From filtered ion charges and types;
- Filter:
  - Precursor charges: 2, 3, 4, 5;
  - Ion charges: 1, 2;
  - Ion types: y, b;

Product ion selection:
From “ion 3”;
To “last ion - 1”;
Special ions:
N-terminal to Proline: checked;
Use DIA precursor window for exclusion: checked;
Auto-select all matching transitions: checked;

(5) Settings -> Peptide Settings:
Modifications:
Structural modifications:
Gln -> pyro-Glu (N-term Q): “Variable” checked;
Pyro-carbamidomethyl (N-term C): “Variable” checked;
Oxidation (M): “Variable” checked;
Glu -> pyro-Glu (N-term E): “Variable” checked;
Carbamyl (N-term) without H: “Variable” checked;
Carbamidomethyl (C): “Variable” unchecked;
Max varable mods: 3;
Max neural losses: 1;
Isotope label type: heavy;
Isotope modifications:
Label: 13C(6)15N(2) (C-term K) checked;
Label: 13C(6)15N(4) (C-term R) checked;
Internal standard type: light;
Filter:
Min length: 7;
Max length: 36;
Exclude N-terminal AAs: 36;
Auto-select all matching peptides: checked;
Prediction:
Retention time predictor: input manually the 10 endogenous precursors in the spectral library without any control preucursors;
Use measured retention times when present: checked;
Time window: 2min;

(6) File -> Import -> transition List.
Skip the warning window;
Add iRT values to the iRT calculator;

(7) Refine -> Add Decoys.
Decoy generation method: Reverse Sequence;

(8) Settings -> Integrate All.

(9) Save the document.

(10) File -> Import -> Results -> Add single-injection replicates in files -> OK.

(11) Refine -> Reintegrate:
Peak scoring model: Add;
Choose model: mProphet;

Use decoys: checked;
Check all of the available feature scores;
Train;
OK;
Integrate all peaks;
Overwrite manual integration: checked;
OK;

(12) File -> Export -> Report.
The report template (SWATHbenchmark_long.skr) provided by LFQbench [2] was used.

(13) Options that are not mentioned above were ignored and their default settings were used.

##### 3. Datasets used in this work

| Dataset | N runs | Equipment | Year | Dataset ID | application |
|---|---|---|---|---|---|
| L929 mouse dataset | 3 | TripleTOF 5600 | 2020 | PXD021390 | Training |
| HEK 293 dataset | 3 | Orbitrap Fusion Lumos | 2020 | PXD015098 | Training |
| SGS human dataset | 30 | TripleTOF5600 | 2014 | PASS00289 | Testing |
| Mouse cerebellum dataset | 10 | Orbitrap Fusion Lumos | 2020 | PXD011691 | Testing |
| LFQbench 64var TTOF6600 dataset | 6 | TripleTOF 6600 | 2018 | PXD002952 | Testing |
| Roland et al. dataset | 4 | Q Exactive HF | 2017 | PXD005573 | Testing |
